# Activity-based selection for enhanced base editor mutational scanning

**DOI:** 10.1101/2024.11.12.622254

**Authors:** Eleanor G Kaplan, Ryan J Steger, Spencer T Shah, Laura M Drepanos, Audrey L Griffith, Ganna Reint, John G Doench

## Abstract

Base editing is a CRISPR-based technology that enables high-throughput, nucleotide-level functional interrogation of the genome, which is essential for understanding the genetic basis of human disease and informing therapeutic development. Base editing screens have emerged as a powerful experimental approach, yet significant cell-to-cell variability in editing efficiency introduces noise that may obscure meaningful results. Here, we develop a co-selection method that enriches for cells with high base editing activity, substantially increasing editing efficiency at a target locus. We evaluate this activity-based selection method against a traditional screening approach by tiling guide RNAs across *TP53*, demonstrating its enhanced capacity to pinpoint specific mutations and protein regions of functional importance. We anticipate that this modular selection method will enhance the resolution of base editing screens across many applications.

## INTRODUCTION

The functional characterization of genetic elements is one of the core challenges of modern biology, leading to the development of numerous experimental approaches. Deep mutational scanning (DMS) relies on overexpression of a library of exogenous open reading frames containing all variants of a gene of interest, allowing for precise mutational interrogation^1,2^. DMS, however, often examines genes outside of their native context and does not account for natural transcriptional and splicing regulation. This approach is therefore often limited by the need for cell line engineering to remove endogenous gene expression^3^. Likewise, massively parallel reporter assays (MPRAs), used to interrogate variants of non-coding regions, also remove genetic elements from their endogenous context^4–6^. Saturation genome editing (SGE) addresses this concern by enabling precise endogenous mutations via homology-directed repair with oligonucleotide templates containing the edit of interest^7,8^. Efficiency limitations, however, have generally required haploid cell models, which limits the generalizability and scalability of this technology. More recently, prime editing has been developed to install a range of precise mutations – insertions, deletions, or single-base pair changes – but currently operates at low efficiency, limiting its use in high-throughput screens to small, contiguous regions of the genome^9–12^.

Base editing offers a high-throughput, endogenous alternative in which an adenine or cytosine deaminase is fused to a Cas protein, enabling A>G or C>T editing within a window of approximately four nucleotides^13–15^. Base editing has been used for a variety of genome scanning purposes, including interrogating drug resistance mechanisms, post-translational modifications, and non-coding regulatory elements^16–22^. Base editing screens, however, are currently hindered by variation in editing activity across a population of cells. Indeed, in large-scale screens, base editors are frequently transduced into cells via lentivirus, resulting in a semi-random locus of integration that may impact editing activity depending on the genomic context^23,24^. Additional factors such as transgene silencing or the cellular immune response to Cas9 may also contribute to this heterogeneity^25,26^. Regardless of the underlying cause, this variation in activity adds noise to screens given that some cells edit highly efficiently and others not at all. Traditional indirect selection methods fail to account for this variation, limiting the resolution of base editing screens and their ability to distinguish biological signal. Here, we develop an activity-based co-selection method to improve the resolution of base editing screens.

## RESULTS

### NG-Cas9 base editor tiling of TP53

To benchmark the high-throughput mutational scanning ability of base editing, we conducted a tiling screen to identify loss-of-function mutations along the highly-studied gene *TP53*^27–31^. Encoding the tumor suppressor protein p53, *TP53* serves a variety of growth control functions, including DNA damage repair and cell cycle regulation via apoptosis^32^. The A549 lung cancer cell line enables both positive and negative selection screens for p53 activity with Nutlin-3 and etoposide, respectively. Nutlin acts as an inhibitor of MDM2, leading to the stabilization of p53 and enrichment of p53 loss-of-function mutations under normal growth conditions^33–35^. Conversely, etoposide induces DNA damage such that loss-of-function mutations deplete given the decreased ability of p53 to induce DNA damage repair^34,36,37^.

We designed a base editing library to include 923 single guide RNAs (sgRNAs) targeting *TP53*, 75 non-targeting sgRNAs, 75 intergenic sgRNAs, and 32 sgRNAs targeting splice donors of pan-lethal genes to serve as positive controls, all using NGN PAMs or known-active PAMs with NG-Cas9^38^. These libraries were cloned into all-in-one base editor lentiviral vectors that contain the PAM-flexible variant NG-Cas9 fused to either the adenine base editor (ABE) or cytosine base editor (CBE) machinery, using the ABE8e and rat APOBEC deaminases, respectively; a 2A peptide confers puromycin resistance (*Fig. 1a*)^15,39,40^. We transduced the sgRNA tiling library into A549 cells in duplicate and selected with puromycin. After selection was complete on day 7, we split the cells into three arms per base editor: no drug, Nutlin, and etoposide. 21 days post-transduction, we retrieved the integrated sgRNA library via PCR of the genomic DNA.

**Figure 1:**
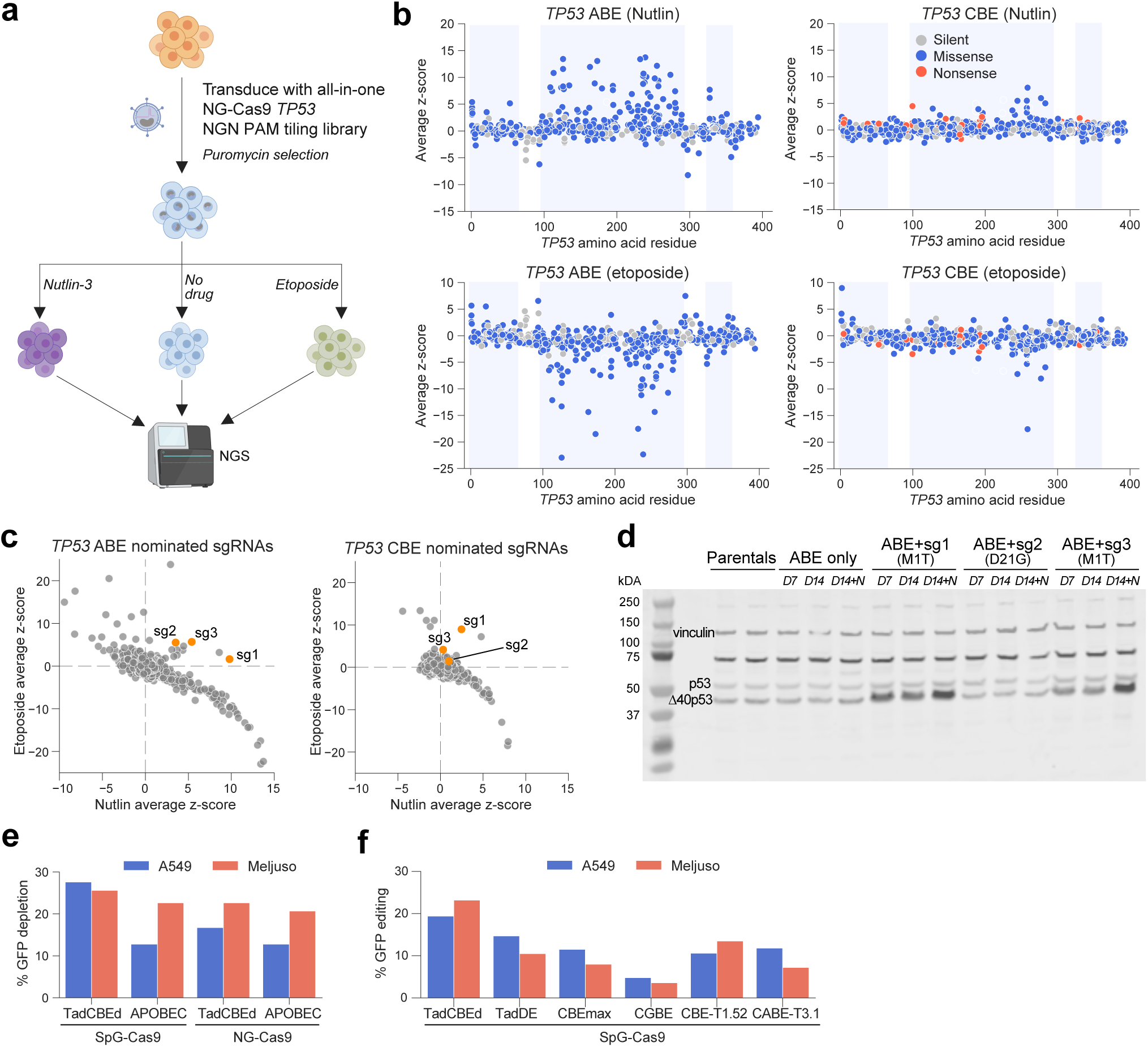
NG-Cas9 base editor tiling of TP53 and CBE optimization (a) Schematic of NG-Cas9 base editor tiling screens, with either ABE8e (ABE) or rat APOBEC (CBE) deaminases. (b) Mean sgRNA z-scores of drug arms relative to no drug arms along the coding sequence of TP53 ABEs (left) and CBEs (right). Domain regions are highlighted in light blue: Left, transactivation domain; Middle, DNA binding domain; Right, oligomerization domain. Color indicates the predicted editing outcome. (c) Scatterplot depicting sgRNA z-scores in etoposide versus Nutlin arms for ABE (left) and CBE (right). sgRNAs nominated for validation are highlighted in orange.(d) Western blot validation of sgRNA 1, 2, and 3, showing p53 versus Δ40p53 levels. Vinculin used as loading control. D=day sample was taken, N=Nutlin. The band at∼75kDa is a known non-specific band of the anti-p53 antibody.(e) GFP activity assay evaluating activity differences between SpG-Cas9 and NG-Cas9 base editors in both A549 and MelJuSo cells as observed by flow cytometry.(f) GFP activity assay benchmarking CBEs with SpG-Cas9 in both A549 and MelJuSo cells as determined by Sanger sequencing and analysis with EditR.

After sequencing deconvolution, we calculated the log2 fold change (LFC) from the pDNA for the no drug (dropout) arm, and for Nutlin and etoposide from the no drug arm. Replicate-level z-scores were calculated relative to the intergenic control sgRNAs and replicates were averaged (*Fig. 1b, Supp. 1a*). As expected, much of the observed signal fell within known *TP53* domains, especially the DNA binding domain (*Supp. 1b*). Notably, in both drug conditions, ABE showed greater magnitude of signal than CBE. Furthermore, most sgRNAs were negatively correlated across the two conditions, enriching with Nutlin and depleting with etoposide, although some sgRNAs deviated from this pattern (*Fig. 1c*). We selected 3 such sgRNAs for validation: sgRNAs 1 and 3, predicted to generate a start codon Met1Thr mutation with ABE, and sgRNA 2, predicted to generate an Asp21Gly mutation with ABE.

We individually delivered these sgRNAs into A549 cells expressing NG-Cas9-ABE8e, selected on puromycin, then split between no drug and Nutlin arms. We verified the identity of the mutations introduced by these sgRNAs by PCR and sequencing of the target locus, analyzing the results with CRISPResso2 software^41^ *(Supp. 1c)*. Previous work has shown that utilization of an alternative downstream start codon at amino acid 40 creates a truncated isoform, Δ40p53, that cannot bind MDM2 and that lacks most of the TAD, which would be consistent with the phenotype observed in the screen^42,43^. To explore the effect of these base edits at the protein level, we performed a Western blot with samples taken at day 7 and day 14 post-transduction (*Fig. 1d*). We indeed observed an increase in a shorter p53 isoform for sgRNAs 1 and 3, but not sgRNA 2, which is consistent with the hypothesis that mutations at the canonical start codon result in the use of an alternative start site.

### CBE optimization

Given the relatively poor performance of the APOBEC-based CBE compared with the ABE in the prior screen, we asked whether alternative Cas9 or CBE variants could increase editing efficiency. Of particular interest were CBEs evolved from ABEs, such as TadCBEd, which have been shown to have higher on-target activity and lower off-target activity than APOBEC^15^. Using the Fragmid vector assembly system^44^, we evaluated the performance of the PAM-flexible Cas9 variants NG-Cas9 and SpG-Cas9^45^ with APOBEC and TadCBEd in A549 and MelJuSo cells. We conducted an activity assay in which GFP is targeted by an sgRNA that can introduce two stop codons into the coding sequence, disrupting fluorescence. Via flow cytometry performed 14 days post-transduction, we observed the highest GFP depletion with SpG-Cas9-TadCBEd across both cell types (*Fig. 1e*). With both CBEs tested, SpG-Cas9 resulted in greater GFP disruption than NG-Cas9, and we therefore proceeded with SpG-Cas9 as our PAM-flexible Cas variant.

With SpG-Cas9, we then benchmarked the performance of three C>T editors (TadCBEd, CBEmax, CBE-T1.52), two dual C>T and A>G editors (TadDE, CABE-T3.1), and one C>G editor (CGBE) (*Fig. 1f*)^15,46–48^; given that TadCBEd outperformed APOBEC in the previous experiment, we did not include APOBEC in this second round of experimentation. With the same GFP activity assay construct, we consistently observed the highest editing efficiency with TadCBEd, with approximately 20% editing in A549 cells and 30% editing in MelJuSo cells, as determined via Sanger sequencing of the target locus and analysis with EditR^49^. We therefore selected SpG-Cas9-TadCBEd as the CBE for future experiments. Even with this more efficient CBE, however, approximately 70% of cells remained unedited in the post-selection population, indicating the need for additional optimization of base editing screening methods.

### Developing a co-selection method to enrich for base editor activity

A standard method for establishing base editor cell models relies on a co-expressed drug resistance or fluorescent marker, an indirect measure of base editor activity hereafter referred to as presence-based selection. This approach does not necessarily result in a high fraction of editing activity as demonstrated above, and the presence of cells in a pooled population that contain an sgRNA but did not edit the target site obscures signal. Inspired by co-selection approaches that enrich for cells with increased homology-directed repair efficiency^50,51^, we set out to develop a co-selection method, referred to as activity-based selection, that directly enriches for cells that have base editing activity at the splice donor site of a selectable marker, rendering the marker functional.

We initially developed this base editing co-selection method by interrupting the puromycin resistance gene with a synthetic intron containing a defective splice donor site (*Fig. 2a*). We derived this sequence from the SV40 intron, which is efficiently spliced in human cells and has been shown to increase transgene expression^52^. As the canonical splice donor site is GT, we designed the 5’ splice site sequence to consist of either AT or GC, which can be edited by ABE or CBE, respectively, to restore splicing. The splice donor sequence corresponds to nucleotides 4 and 5 of the sgRNA, as these positions are highly effective for base editing. The total variable target site is 20-nucleotides long, with 3 nucleotides targeting the exon portion of the puromycin resistance gene and the other 17 targeting the intron, followed by an NGG PAM (*Fig. 2b*). In order to maximize the likelihood of splicing upon successful editing, variations of the SV40 intron were designed using common motifs derived from a sequence consensus collated from over 200,000 mammalian splice sites^53,54^. Further, the intron variants avoid adenines and cytosines in positions 6 - 9 to mitigate the possible confounder of multiple edits. To maximize the likelihood of base editing activity, these intron sequences were chosen such that their targeting sgRNAs have high Rule Set 3 scores as determined by CRISPick^55^. The intron also contains a stop codon such that unspliced transcripts will result in premature termination of translation. With these parameters, we designed two libraries – one for each base editor – each containing 200 intron sequences.

**Figure 2:**
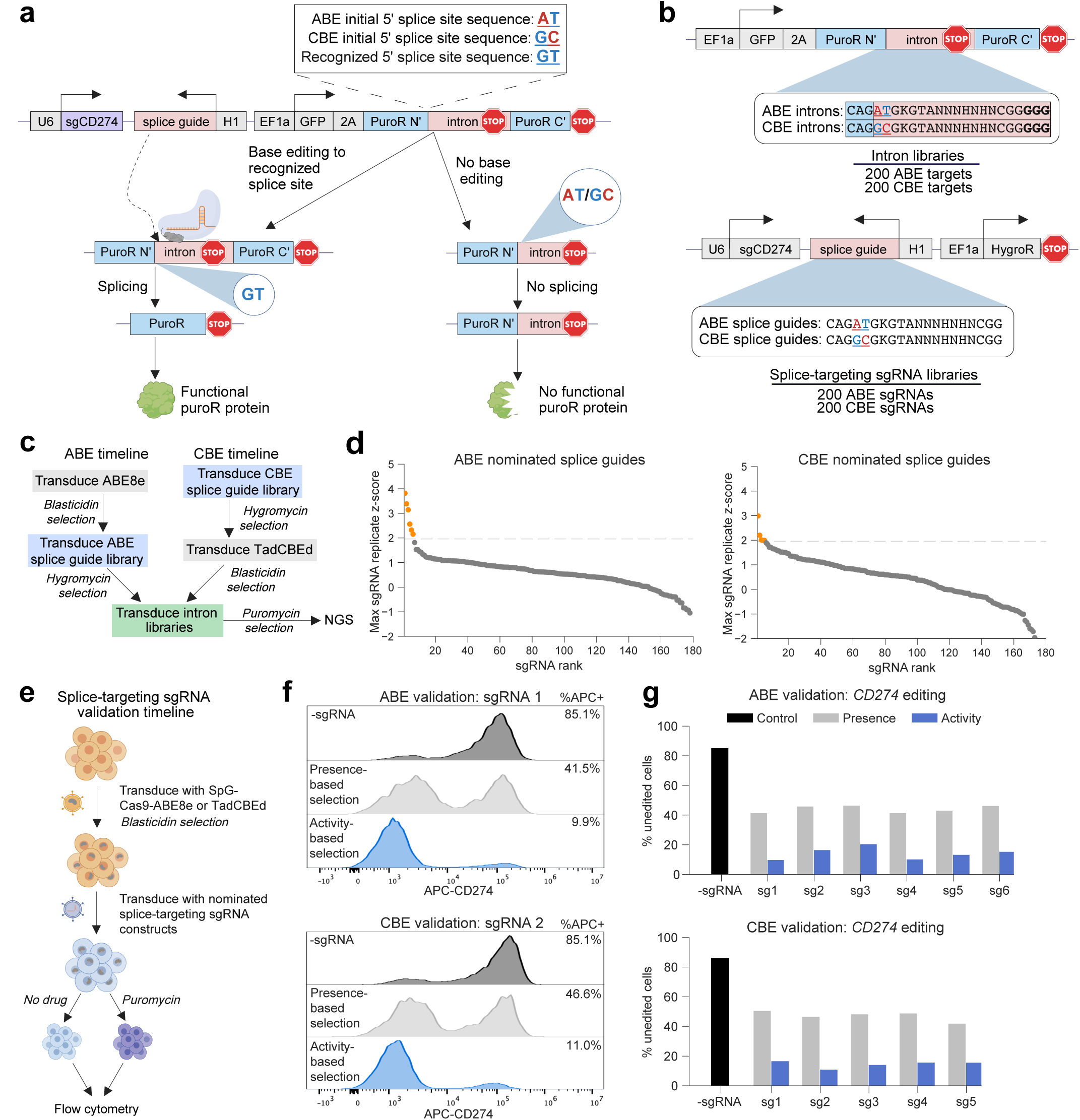
Activity-based selection method development (a) Schematic depicting base editing activity-based selection screen design. Base editors can restore the intron splice donor sequence to the recognized mammalian consensus sequence, allowing for splicing and functional protein expression. If no editing occurs, no splicing occurs and no functional protein is expressed. (b) Vector schematics showing variation in introns (top) and corresponding splice-targeting sgRNAs (bottom). Nucleotides in red can be base edited to the splice donor consensus sequence (GT). Nucleotides in blue are part of the consensus sequence. The blue box depicts nucleotides within the puromycin resistance exon. The pink box depicts nucleotides within the intron. PAM is shown in bold. (c) Timelines of ABE (left) and CBE (right) primary screens. Screens followed the same timeline beginning with the intron library transduction. (d) Scatter-plot of maximum sgRNA z-scores for ABE (left) and CBE (right) screens. Horizontal dashed line represents validation cutoff of z>=1.96. Dots in orange represent sgRNAs nominated for validation. BE = base editor corresponding to splice-targeting sgRNA used. (e) Schematic of splice-targeting sgRNAs validation timeline. (f) Example of splice-targeting sgRNA validation results for ABE (top) and CBE (bottom). CD274 editing was measured by flow cytometry. (g) Validation of all nominated primary screen splice-targeting sgRNA. Bars depict the percent of unedited cells remaining with each selection method for ABE (top) and CBE (bottom) modalities.

Accordingly, we designed two libraries – one for each base editor – each containing 200 sgRNAs that target the corresponding intron sequence at the 5’ splice site of the puromycin resistance gene. We cloned these two splice-site-targeting sgRNA libraries into vectors that also include an sgRNA targeting the wildtype 5’ splice site of the nonessential cell-surface marker *CD274* (gene ID: PD-L1), in order to assess if base editing activity at one locus corresponds to base editing activity at a distinct endogenous locus (*Fig. 2b*). When base editing occurs on the antisense strand of *CD274*, the 5’ splice site is disrupted and PD-L1 expression decreases.

We screened each combination of splice-site-targeting sgRNA library and corresponding intron variant library in duplicate. For the ABE arm, we first transduced ABE8e into A549 cells and selected with blasticidin, then transduced the splice-site-targeting sgRNA library and selected with hygromycin. For the CBE arm, we first transduced the splice-site-targeting sgRNA library and selected with hygromycin, then transduced TadCBEd and selected with blasticidin. Lastly for both arms, we transduced the corresponding intron library, selected with puromycin, and at the end of the screen used PCR to retrieve the enriched splice-site-targeting sgRNAs (*Fig. 2c, Supp. 2a*). Replicates were more correlated for CBE (r=0.65-0.95) than ABE (r=0.2-0.41). We attribute this difference to the differing timelines, as the ABE was removed from selective pressure for three weeks longer than the CBE, possibly allowing for an increase in the heterogeneity of ABE expression.

### Splice-targeting sgRNA validation

To validate the screen and identify top-performing sgRNA - intron pairs, we nominated all sgRNAs with at least one replicate z-score greater than or equal to 1.96, resulting in 6 ABE and 5 CBE sgRNAs (*Fig. 2d*). To minimize the chance of off-target activity, we ensured all nominated sgRNAs did not map to a perfect or 1-nucleotide mismatch from another sequence in the genome by using Cas-OFFinder^56^. We assembled 11 new constructs, each containing a unique splice-targeting sgRNA and its corresponding intron sequence disrupting the puromycin resistance gene, as well as an intact, upstream GFP marker. To evaluate the generalizability of this approach, we performed this validation experiment in MelJuSo cells.

After introducing ABEs or CBEs into cells and selecting with blasticidin, we separately transduced the 11 sgRNA constructs and divided cells between no drug (presence-based selection) and puromycin (activity-based selection) conditions 4 days post-transduction. Sixteen days post-transduction, we performed flow cytometry to assess CD274 protein levels (*Fig. 2e*). For both selection conditions, we gated for expression of GFP to identify cells transduced with the sgRNA. When comparing presence-versus activity-based selection methods, we observed that activity-based selection decreased the number of unedited cells in the final selected population from 41.5% to 9.9% with the most active ABE sgRNA, and from 46.6% to 11.0% with the most active CBE sgRNA (*Fig. 2f*). Across both base editors, activity-based selection resulted in a substantial decrease in the number of unedited cells in the final population for every sgRNA - intron pair nominated, indicating the robustness of this approach (*Fig. 2g*). We confirmed the levels of editing of the activity-based selection condition by sequencing the endogenous *CD274* locus and analyzing per-base editing efficiency with the EditR analysis tool^49^ (*Supp. 2b*). We saw greatly increased editing at C6 compared to C5, likely due to the lower likelihood of editing observed with TadCBEd when the cytosine follows an adenine^15^. We selected the two sgRNA - intron pairs with the highest activity-based selection editing efficiency from each base editor arm (ABE: sgRNA 1, 4; CBE: sgRNA 2, 3) for further evaluation.

The modular design of the co-selection system allows for easy adaptation to other selectable markers given that 17 out of 20 nucleotides targeted by the splice sgRNA lie within the intron sequence, anchored to the exon by just three nucleotides, CAG. To demonstrate this flexibility, we inserted the intron within the hygromycin resistance gene and observed similarly-improved performance of activity-based selection (*Supp. 2c*). Additionally, we inserted the intron into GFP to evaluate if cryptic splicing is occurring in the system, which would bypass the need for base editors. We transduced this construct into cells with and without base editors and observed minimal GFP positivity (<1%) in cells without base editors, indicating a requirement for base editing activity (*Supp. 2d*).

We next assessed if activity-based selection could also increase the editing efficiency of SpRY-Cas9, a nearly PAM-less Cas9 variant^45,57^. For both ABE and CBE, we transduced two validated sgRNA vectors into SpRY-Cas9-base editor-expressing cells, each with their corresponding intron sequence disrupting the puromycin resistance gene, and then evaluated *CD274* editing via flow cytometry. Most cells (>80%) remained unedited after selection, and we noticed no improvement with the activity-based selection approach (*Supp. 2e*). We hypothesized that this low efficiency editing may be due to self-editing of the sgRNA, as previously observed^58^. Due to its lack of a PAM requirement, SpRY-Cas9 sgRNAs may recognize and edit the integrated guide cassette, creating a mutated guide that can no longer recognize its target sequence, thus resulting in decreased editing efficiency. To test this hypothesis, we performed Sanger sequencing of the integrated guide cassette targeting *CD274*. For comparison, we did the same for cells with SpG-Cas9, which should not be able to self-edit given PAM restrictions. For all samples, the most edited nucleotides fell within the canonical base editing window of 4-8 nucleotides along the integrated guide. SpRY-Cas9 showed the highest self-editing at positions 4 and 6, with 87.5% A>G editing at nucleotide 4 along the guide and 86.0% editing at nucleotide 6 (*Supp. 2f*). Surprisingly, cells with SpG-Cas9 also showed detectable editing of the integrated sgRNA (editing at A4: 28.5%, C6: 38.0%). Given the high rate of sgRNA mutations, we believe that self-editing inhibits the use of activity-based selection with SpRY-Cas9 under these experimental conditions.

### Pooled screening with TP53

Given the success of SpG-Cas9 base editor co-selection at increasing editing efficiency at a single locus, we next asked if this approach could generalize across a library of sgRNAs. We opted to re-screen *TP53* to directly compare this selection approach with traditional presence-based selection (*Fig. 3a)*. We designed a new library to include 2,785 sgRNAs tiling along the length of *TP53* with no PAM restriction (i.e. NNN PAM; we had been hopeful that SpRY-Cas9 would prove effective), 206 intergenic controls, and 103 pan-lethal splice site positive controls, which utilize an NGG PAM. We cloned this library into two types of vectors: a presence-based selection vector with an intact puromycin resistance gene and an activity-based selection vector with an intron-disrupted puromycin resistance gene and its corresponding splice-targeting sgRNA (*Fig. 3b*). To conduct this screen, we first established A549 cells containing SpG-Cas9-ABE8e or -TadCBEd on blasticidin. We then transduced the *TP53* tiling library and selected on puromycin starting on day 2 for the presence-based selection version and day 4 for the activity-based selection versions, giving extra time for the latter to allow for editing to occur at the puromycin resistance intron. Following puromycin selection, cells were split between etoposide and no drug conditions; we opted for a negative-selection condition because data quality of such screens is more reliant on high editing activity. After two weeks of additional passaging, we harvested cells and sequenced the sgRNA library.

**Figure 3:**
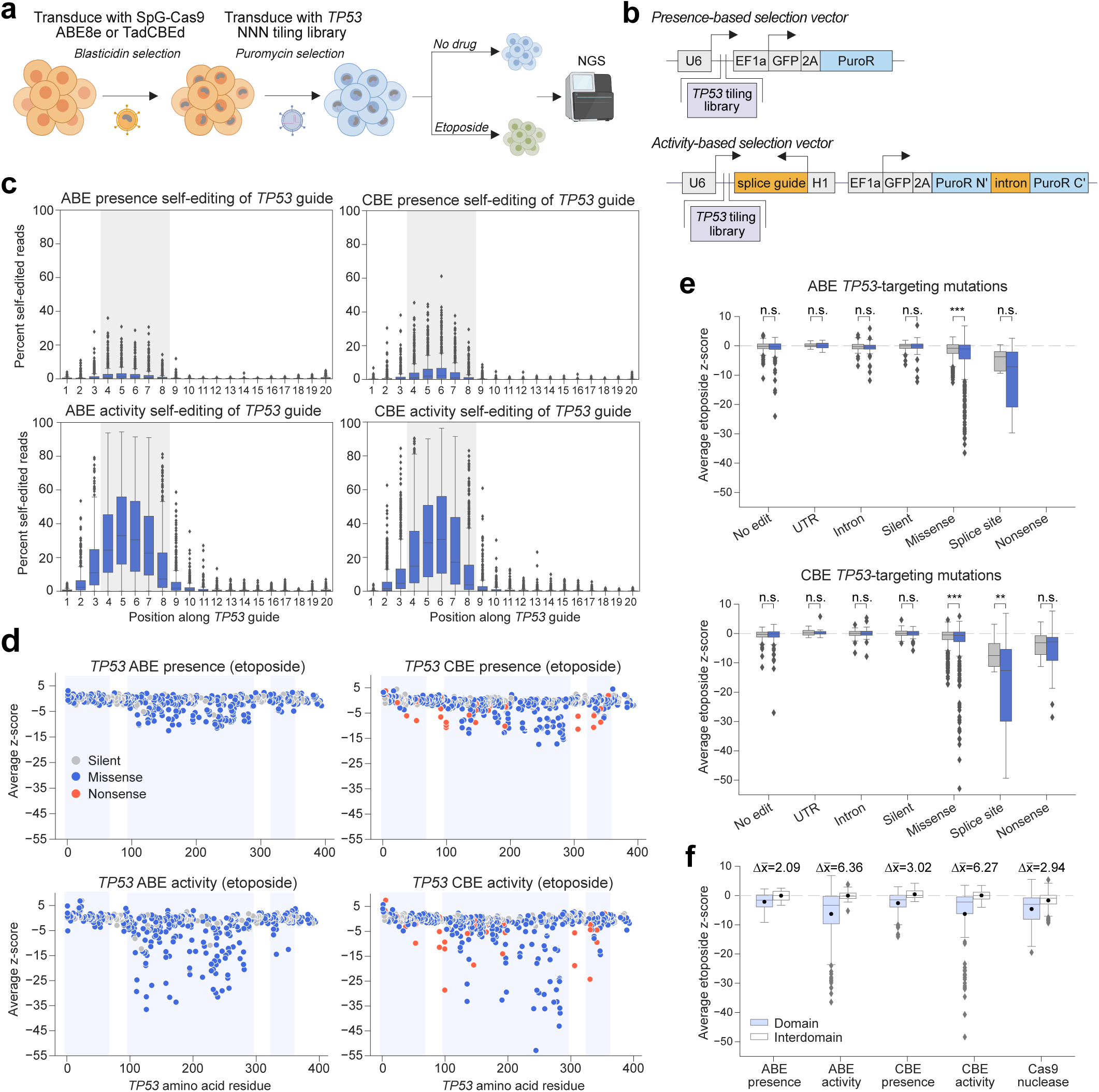
Activity-based selection base editor tiling of TP53 (a) Schematic of SpG-Cas9 base editor tiling screen.(b) Depiction of selection method constructs. Presence-based selection construct (top) contains an intact puromycin resistance gene. Activity-based selection construct (bottom) contains an intron-disrupted puromycin resistance gene and a previously-validated splice-targeting sgRNA.(c) Self-editing analysis: percent edited reads by position along TP53 sgRNA, including both drug and no drug arms. The canonical editing window of nucleotides 4-8 is highlighted in gray. Boxes depict 25th and 75th percentiles as minima and maxima and the center represents the median; whiskers depict 10th and 90th percentile. (d) Average etoposide z-scores relative to no drug arm by base editor and selection condition. sgRNAs are colored by predicted mutation and plotted by median targeted residue. Only sgRNAs targeting coding regions of TP53 are depicted. Domain regions are highlighted in light blue. (e) Average etoposide z-scores for all ABE (top) and CBE (bottom) TP53-targeting mutations separated by predicted mutation. Boxes depict 25th and 75th percentiles as minima and maxima and the center represents the median; whiskers depict 10th and 90th percentile. Stars depict p-value significance according to the following: p-value <= 0.001: ***, <= 0.01: **, <= 0.05: *, >0.05: n.s. (two-sided t-test). (f) Difference in means between z-scores within annotated domain versus interdomain regions of TP53 by selection method and technology. Domain regions are outlined in Supplementary Figure 1b. Boxes depict 25th and 75th percentiles as minima and maxima and the center represents the median; whiskers depict 10th and 90th percentile. Black dots represent the mean.

Upon sequencing deconvolution, we observed an approximately 20% lower match rate between the observed and expected sgRNA sequences with activity-based selection than with presence-based selection (*Supp. 3a*), which we attributed to self-editing, as observed above with SpG-Cas9. This may be possible if the sgRNA forms a one-nucleotide bulge to recognize the first guanine of the tracrRNA sequence as the middle guanine of the NGN PAM used by SpG-Cas9 (*Supp. 3b).* Alternatively, GTT, the beginning of the tracrRNA sequence, may serve as a PAM for SpG-Cas9 under these conditions. Regardless of the mechanism, to further explore this phenomenon, we obtained the top one million unexpected sequencing reads and mapped them back to their original sgRNAs given any A>G or C>T editing. We then calculated the percent of unexpected reads out of the total reads for each sgRNA and observed that activity-based selection resulted in substantially higher self-editing rates overall (*Supp. 3c*). For each sgRNA, we also calculated the percent of base-edited nucleotides (G or T) out of the total reads for that nucleotide across all screening conditions. With both selection methods, the most-edited bases fell within the expected base editing window of 4 - 8 nucleotides along the sgRNA (*Fig. 3c*), further supporting the self-editing theory. We observed much lower rates of editing in the presence-based selection condition, indicating that typical selection methods do not result in substantial self-editing, which likely explains why this phenomenon has not, to our knowledge, been previously reported.

To account for this self-editing, we applied a computational correction to add the read counts from the unexpected sequences to the read counts of the original sgRNAs to which they mapped. We then calculated the LFC of the no drug arms relative to the pDNA, and the LFC of the etoposide arms relative to the no drug arms. We observed that the computational correction improved replicate correlations (*Supp. 3d*), with the greatest increase occurring under the activity-based selection conditions, as expected given their higher rates of self-editing. We therefore proceeded with the analysis using the corrected LFC values.

For sgRNAs targeting *TP53*, we restricted the subsequent analysis to sgRNAs with an NGN PAM given that we screened this library with SpG-Cas9. Notably, the positive control pan-lethal-targeting sgRNAs showed a wider dynamic range of LFC values with activity-based selection compared to presence-based selection, while the negative control intergenic sgRNA distribution remained similar between methods (*Supp. 3e*). We then z-scored LFC values for sgRNAs targeting *TP53,* relative to the intergenic controls, and saw a substantially wider dynamic range of z-scores with activity-based selection relative to presence-based selection (*Fig. 3d, Supp. 3f*). Whereas the z-scores in the presence-based selection conditions range from -12.5 to 4.2 and -17.4 to 8.4 for ABE and CBE, respectively, activity-based selection showed a wider range of -36.5 to 15.9 for ABE and -52.9 to 17.7 for CBE. When discretized by predicted mutation bin, we observed that sgRNAs predicted to introduce either a silent edit or no edit showed no significant difference between the selection regimes for both ABE and CBE, whereas missense mutations were significantly more depleted with activity-based selection compared to presence-based selection (*Fig. 3e*), indicating that activity-based selection did not indiscriminately increase the range of the screen but rather substantially increased the ratio of signal-to-noise.

Lastly, we assessed the ability of base editing, in comparison to CRISPR knockout – another widely used method for endogenous high-throughput interrogation – to identify known functional domains of *TP53*^59–63^. With CRISPR knockout, for every insertion or deletion created, there is only, on average, a ⅓ chance that the repair outcome results in an in-frame mutation. Out-of-frame mutations, regardless of whether they fall within functional domains, will likely lead to a non-functional protein, and thus the overall resolution of this approach is limited by the need to distinguish the signal produced by ⅓ of mutations from the noise produced by the other ⅔. To compare the performance of these approaches for identifying domains, we screened the same *TP53* library in cells expressing SpG-Cas9-nuclease. We observed the greatest difference in mean z-score between domain versus interdomain regions for the base editor activity-based selection methods (ABE: x̄ = 6.36; CBE: x̄ = 6.27) (*Fig. 3f*). Comparatively, presence-based selection methods showed less distinction between domain and interdomain regions (ABE: x̄ = 2.09; CBE: x̄ = 3.02). Cas9-nuclease showed a similar distinction as the presence-based selection method between domain and interdomain regions (x̄ = 2.94), but also showed depletion within interdomain regions (x = -1.68), likely due to frameshift mutations in interdomain regions disrupting the overall protein structure. Conversely, for base editors across all selection methods, the means of interdomain regions centered near 0 (ABE presence: x = -0.06; CBE presence: x = 0.42; ABE activity: x = -0.05; CBE activity: x = -0.03). Overall, base editing outperformed Cas9-nuclease for distinguishing domain versus interdomain regions, and activity-based selection more clearly delineated these functional regions compared to traditional selection methods.

## DISCUSSION

BE screens offer a scalable approach to interrogating endogenous genetic elements, but variation in editing activity on a cell-to-cell basis limits screen resolution. Here, we develop a splice-based selection method to enrich for actively editing cells, reducing the number of unedited cells from over 40% to less than 10% at a single locus. Additionally, we demonstrate increased signal-to-noise ratio in a pooled, negative selection screen tiling *TP53*, identifying specific point mutations and domains of functional importance.

Given the modularity of this splice-targeting sgRNA and intron system, activity-based selection can be seamlessly incorporated into most base editing experiments to more clearly detect potential gain- and loss-of-function mutations. Beyond drug selection, for example, the intron could be inserted within a fluorescent reporter such that actively-editing cells could be enriched for using fluorescence-activated cell sorting. This approach could be especially useful for models that rely on transient base editor delivery, such as via protein or mRNA, as such non-integrating delivery methods currently have no straightforward means of selecting for cells that received sufficient levels of the base editor^19,64^. Activity-based selection mitigates this transient delivery challenge for base editors by generating a stable selection marker. Finally, simply by altering the PAM sequence in the synthetic intron, this approach could also be repurposed for use with other Cas proteins.

One limitation of activity-based selection when using the SpG variant of Cas9 is the increased propensity for self-editing. We have developed a computational approach to rescue the sequencing read counts from self-edited sgRNAs, although we note that this does not account for the subsequent expanded range of potential off-target sites for the mutated sgRNA. Further, while more apparent with activity-based selection, self-editing activity was also observed with traditional selection, underscoring the need to correct for this phenomenon when screening with PAM-flexible Cas9 varieties, lest the cells with the most-active base editing activity fall out of the analysis due to the sgRNA no longer being recognized during sequencing deconvolution. The potential for self-editing, alongside the ability of base editors to generate multiple alleles, highlights the need for subsequent validation of the editing outcome at the target site.

We envision that the increased sensitivity of this selection method will enhance the ability of base editors to uncover the functional roles of poorly characterized genomic regions, including underexplored genes and non-coding areas with regulatory potential. Base editing screens are easily applied to models that have been proven scalable for genome-wide interrogation by other CRISPR techniques, such as knockout, interference, or activation, enabling the investigation of tens to hundreds of thousands of sgRNAs. Further, screens focused on a specific gene or gene set can uncover drug resistance mechanisms, map functional domains, and identify rare gain-of-function mutations^18,20,65–67^. Following a base editing screen, genomic regions of interest can be further refined with more precise editing technologies that are best deployed at focused loci, such as saturation genome editing or prime editing. Together, this suite of tools offers great potential for understanding disease mechanisms and informing drug discovery efforts.

## ACKNOWLEDGEMENTS

We thank all members of the Genetic Perturbation Platform (GPP) production teams: Desiree Hernandez, Monica Roberson, Berta Escude Velasco, Eliezer Josue Ibarra, Noah Smith, Dylan McLaughlin, Tashi Lokyitsang, and Xiaoping Yang for producing sgRNA libraries and lentivirus; Abrianna Bowie, Olivia Bare, Yenarae Lee, Quinton Celuzza, Meredith Felt, and Amy Herman for logistics support; Matthew Greene, Doug Alan, Mark Tomko, Bronte Wen, Angeline Tamayo, and Tom Green for software engineering support; Dave Root for GPP leadership; the Broad Institute Genomics Platform Walk-up Sequencing group for Illumina sequencing; the Functional Genomics Consortium for funding support. We especially thank Nathan Miller and Ify Nwolah for laboratory assistance, Fengyi Zheng for programming advice, Allison Uebele for helpful discussions, and Sarra Merzouk for a close reading of the manuscript.

## METHODS

### Table of Reagents

| Reagent or Resource | Source | Identifier |
| --- | --- | --- |
| **Antibodies** |  |  |
| Mouse Anti-p53 Monoclonal Antibody | Abcam | Cat#ab28, RRID:AB_303312 |
| Recombinant Anti-Vinculin antibody | Abcam | Cat#ab219649, RRID:AB_2819348 |
| IRDye 800CW Goat anti-Mouse IgG | LI-COR Biosciences | Cat#925-32210, RRID:AB_2687825 |
| IRDye 680RD Goat anti-Rabbit IgG | LI-COR Biosciences | Cat#925-68071, RRID:AB_2721181 |
| APC anti-human CD274 | Biolegend | Cat#393610, RRID: AB_2749927 |
| **Chemicals** |  |  |
| Nutlin-3 | Selleckchem | Cat#S1061 |
| Etoposide | Sigma-Aldrich | Cat#E1383 |
| Puromycin | Sigma-Aldrich | Cat#P9620 |
| Blasticidin | Thermo Fisher | Cat#A1113903 |
| Hygromycin | Thermo Fisher | Cat#10687010 |
| Polybrene | Sigma-Aldrich | Cat#H9268 |
| TransIT-LT1 | Mirus | Cat#2306 |
| Interferon-γ | Thermo Fisher | Cat#300-02-100UG |
| **Experimental models: Cell lines** |  |  |
| Human A549 cells | Cancer Cell Line Encyclopedia | N/A |
| Human MelJuSo cells | Cancer Cell Line Encyclopedia | N/A |
| Human HEK293T cells | ATCC | ATCC CRL-3216 |
| **Recombinant DNA** |  |  |
| Plasmid pRDA_867 | Addgene | Depositing with Addgene |
| Plasmid pRDB_092 | Addgene | Depositing with Addgene |
| Plasmid pRDB_270 | Addgene | Depositing with Addgene |
| Plasmid pRDB_345 (ABE splice cloning) | Addgene | Depositing with Addgene |
| Plasmid pRDB_347 (CBE splice cloning) | Addgene | Depositing with Addgene |
| Plasmid pRDB_268 | Addgene | Depositing with Addgene |
| **Software and algorithms** |  |  |
| PoolQ | The Broad Institute | https://portals.broadinstitute.org/gpp/public/software/poolq |
| CRISPick | The Broad Institute | https://portals.broadinstitute.org/gppx/crispick/public |
| Fragmid | The Broad Institute | https://portals.broadinstitute.org/gppx/fragmid/public |
| EditR | University of Minnesota | http://baseeditr.com/ |
| CRISPResso2 | Massachusetts General Hospital | http://crispresso2.pinellolab.org |

### Tiling library designs

sgRNA sequences for NG-Cas9 (expanded NGN PAM) and SpG-Cas9 (NNN PAM) *TP53* tiling libraries were designed using *TP53* Ensembl transcript ENST00000269305.8. We included all sgRNAs targeting coding sequences as well as all sgRNAs for which the start was up to 50 nucleotides into the intron and UTRs. NGN PAM libraries included most NGN PAMs as well as other designated active PAMs observed with NG-Cas9^38^. We filtered out sgRNAs that created BsmBI restriction sites or sgRNAs that had more than three consecutive T’s.

### Base editor tiling library annotation

All adenines or cytosines in positions 4-8 of the sgRNA (where 1 is the most PAM-distal position, and positions 21-23 are the PAM), were considered to be edited for ABE and CBE conditions, respectively. The assigned amino acid residue for an sgRNA, regardless of the predicted edit, was calculated based on the position of nucleotides 4-8 of each sgRNA within the coding sequence. Nucleotide edits were used to predict amino acid changes of each sgRNA. Mutation bins were designated by predicted amino acid mutation type. sgRNAs containing multiple mutation types were binned as the most severe mutation type given the following order: Nonsense > Splice site > Missense > Intron > Silent > UTR. sgRNAs with no adenines of cytosines within the editing window for ABEs or CBEs, respectively, were binned as ‘‘No edits.’’

### TP53 domain annotation

Annotations for the three primary functional domains of *TP53* were aligned with those provided by Gould and colleagues^12^: the N-terminal transactivation domain (TAD) spans amino acids 1–61, the central DNA-binding domain (DBD) encompasses amino acids 96–292, and the C-terminal tetramerization domain (TD), also known as the oligomerization domain, includes amino acids 324–356.

### Splice-targeting sgRNA library design

Splice-targeting sgRNAs that target the disrupted intron splice site were designed following the consensus sequence CAG**A**TGKGTANNNHNHNCGG (ABE) or CAGG**C**GKGTANNNHNHNCGG (CBE), with the bold letter representing the base edit necessary to occur for spliceosome recognition. Nucleotides 1-10 are derived from the human 5’ splice site consensus sequence in order to maximize chances for splicing with the following exceptions: nucleotide 6 was assigned a G and nucleotide 7 was assigned a K such that only one nucleotide is editable within the standard base editing window^54^. Nucleotides 11-20 were designed to balance sequence variability with parameters defined by Rule Set 3 to maximize sgRNA efficiency^55^. Parameters included the following: nucleotides 14 and 16 are anything but G, nucleotides 18, 19, and 20 are assigned CGG, respectively. sgRNAs that created BsmBI restriction sites or sgRNAs that had more than three consecutive T’s were removed. Five thousand of the remaining sgRNAs were run through CRISPick to characterize on and off-target activity predictions. sgRNAs were filtered for an on-target score >=0.7 and an off-target rank <500. Of the remaining 857 sgRNAs, 200 ABE and 200 CBE sgRNAs were randomly selected to compose the splice-targeting sgRNA libraries.

### Intron design

Synthetic introns were derived from the SV40 intron and inserted after base 310 of the puromycin resistance gene. The primary intron was modified to contain a stop codon beginning at 23 nucleotides from the start of the codon. For the primary screen, intron libraries corresponding to the splice-targeting sgRNA libraries were created for both ABE and CBE conditions. A second intron was adapted from the previous intron to include additional stop codons beginning at 27 and 31 nucleotides from the start of the intron, such that there is a stop codon within every reading frame. All intron sequences corresponding to splice-targeting sgRNAs used after the primary SpG-Cas9 *TP53* screen are listed in Supplementary Table 1. The intron was inserted in the +2 reading frame of the puromycin resistance gene, the +0 reading frame of the hygromycin resistance gene, and the +1 reading frame of GFP.

### Modular vector design

All pF plasmids were made by gene synthesis into the EcoRV site of pUC57-Kan (Genscript).

### Modular vector destination vector pre-digest

Destination vectors were pre-digested using BbsI and NEBuffer 2.0 (New England Biolabs) at 37°C for 2 h. Linearized destination vectors were gel purified using 0.7% agarose gels and extracted with the Monarch DNA Gel Extraction Kit (New England Biolabs), before further purification by isopropanol precipitation.

### Modular vector Golden Gate assembly

The pF vectors were diluted to 10 nM in sterile water and cloned into a pre-digested destination vector via Golden Gate cloning. Each reaction contained 3 μL BbsI (New England Biolabs), 1.25 μL T4 ligase (New England Biolabs), 3 μL of 10x T4 ligase buffer (New England Biolabs), 75 ng destination vector, and a 1:1 molar ratio of fragments:destination vector. Reactions were carried out under the following thermocycler conditions: (1) 37°C for 5 min; (2) 16°C for 5 min; (3) go to (1), x100; (4) 37°C for 30 min; (5) 65°C for 20 min. The Golden Gate product was treated with Exonuclease V (New England Biolabs) at 37°C for 30 min before enzyme inactivation with the addition of EDTA to 11 mM. Per reaction, 10 μL of product was transformed into Stbl3 chemically competent E. coli (Invitrogen) via heat shock, and grown at 37°C for 16 h on agar with 100 μg/mL carbenicillin. Colonies were picked and grown at 37°C for 16 h in 5 mL Luria-Bertani (LB) broth with 100 μg/mL carbenicillin. Plasmid DNA (pDNA) was prepared (QIAprep Spin Miniprep Kit, Qiagen). Purified plasmids were verified by restriction enzyme digest and whole plasmid sequencing through Plasmidsaurus.

### Library production

Oligonucleotide pools for CP1609 and CP1610 were synthesized by TWIST; CP1986, CP1987, CP1988, and CP1989 were synthesized by IDT; pools for CP2087, CP2088, CP2089, CP2090, and CP1845 were synthesized by Genscript. BsmBI recognition sites were appended to each sgRNA sequence along with the appropriate forward and reverse overhang sequences (bold italic) for cloning into the sgRNA expression plasmids, as well as primer sites to allow differential amplification of subsets from the same synthesis pool. For CP1609, CP1610, CP2087, CP2088, CP2089, and CP2090, the final oligonucleotide sequence structure was thus: 5′-[Forward Primer]CGTCTCA***CACC***G[sgRNA, 20 nt]***GTTT***CGAGACG[Reverse Primer]-3’. For CP1986 and CP1987, the sequence structure was the same except the forward and reverse overhang sequences were replaced with the following: ***TCCC***, ***GTTT.*** For CP1988 and CP1989, the forward and reverse overhang sequences were as follows: ***CTGG***, ***GGGT***.

Primers were used to amplify individual subpools using 25 μL 2x NEBnext PCR master mix (New England Biolabs), 2 μL of oligonucleotide pool (∼30-300 ng), 5 μL of primer mix at a final concentration of 0.5 μM, and water to a final volume of 50 µL. PCR cycling conditions: (1) 98°C for 1 min; (2) 98°C for 30 sec; (3) 53°C for 30 s; (3) 72°C for 30 s; (4) go to step 2, x6-14 depending on library size; (5) 72°C for 5 min.

The resulting amplicons were PCR-purified (Qiagen) and cloned into their respective library vector via Golden Gate cloning with Esp3I (Fisher Scientific) and T7 ligase (Epizyme) under the following thermocycler conditions: (1) 37°C for 5 min; (2) 20°C for 5 min; (3) go to step 1, x100; (4) 37°C for 30 min; (5) 65°C for 10 min. The ligated product was isopropanol precipitated and electroporated into Stbl4 electrocompetent cells (Invitrogen) and grown at 37°C for 16 h on agar with 100 μg/mL carbenicillin. Colonies were scraped and plasmid DNA (pDNA) was prepared (HiSpeed Plasmid Maxi, Qiagen). To confirm library representation and distribution, the pDNA was sequenced by Illumina MiSeq.

### Lentivirus production

For small-scale virus production, the following procedure was used: 18 h before transfection, HEK293T cells were seeded in 6-well dishes at a density of 1x10^6^ cells per well in 1 mL of DMEM + 10% heat-inactivated FBS. Transfection was performed using the TransIT-LT1 transfection reagent (Mirus) according to the manufacturer’s protocol. Briefly, for each construct, 324μL of Opti-MEM (Corning) and 17μL LT1 was combined with a DNA mixture of the packaging plasmid pCMV_VSVG (125 ng; Addgene 8454), psPAX2 (1250 ng; Addgene 12260), and the transfer vector (312.5 ng). The solutions were incubated at room temperature for 30 min and added dropwise to cells. Plates were then transferred to a 37°C incubator for 6–8 h, after which the media was removed and replaced with DMEM + 10% FBS media supplemented with 1% BSA. Virus was harvested and filtered 40 hours after this media change.

A larger-scale procedure was used for pooled library production. 18 h before transfection, 18x10^6^ HEK293T cells were seeded in a 175 cm^2^ tissue culture flask and the transfection was performed the same as for small-scale production using 6 mL of Opti-MEM, 305 μL of LT1, and a DNA mixture of pCMV_VSVG (5 μg), psPAX2 (50 μg), and 10 μg of the transfer vector. Following addition of the transfection mix, flasks were transferred to a 37°C incubator for 6–8 h, then the media was aspirated and replaced with BSA-supplemented media; virus was harvested and filtered 40 h after this media change.

### Determination of lentiviral titer

To determine lentiviral titer for transductions, cell lines were transduced in 12-well plates with a range of virus volumes (e.g., 0, 150, 300, 500, and 800 μL virus) with 3e6 cells per well in the presence of polybrene. For transduction, the plates were centrifuged at 821 x g for 2 h, after which 2 mL of warm media was added to reduce viral toxicity. Plates were then transferred to a 37°C incubator for 4–6 h. Each well was then trypsinized and pooled. Two days post-transduction, we seeded an equal number of cells into two wells of a 6-well plate, and added puromycin to one well. The following passage, both wells were counted for viability. A viral dose resulting in ∼30% transduction efficiency, corresponding to an MOI of ∼0.35, was used for subsequent library screening.

### Small molecule dosages

The dosages for the selection drugs puromycin, blasticidin, and hygromycin were as follows for the relevant cell lines:

A549: puromycin 1.5 μg/mL; blasticidin 5 μg/mL; hygromycin N/A. MelJuSo: puromycin 1 μg/mL; blasticidin 4 μg/mL; hygromycin 100 μg/mL.

Puromycin selection was completed over 5-7 days, while blasticidin and hygromycin selection were completed over 12-14 days. For drug screens in A549 cells, etoposide and Nutlin-3 were dosed at 5 µM and 2.5 µM respectively over the course of the experiment.

### Lentiviral transduction to establish stable cell lines

In order to establish stable Cas-expressing cell lines for screens, MelJuSo cells or A549 cells were transduced with pRDA_867, pRDB_092, pRDB_268, or pRDB_270 in the presence of polybrene at a dosage of 4 µg/mL for MelJuSo cells and 1 µg/mL for A549 cells. Cells were centrifuged at 821 x g for 2 h in 12-well plates, after which 2 mL of warm media was added to reduce viral toxicity. Plates were then transferred to a 37°C incubator for 4–6 h. Replicate wells were then trypsinized and pooled. Successfully infected cells were selected for with blasticidin as described above.

### Pooled screens

For pooled screens, cells were transduced in 2 biological replicates with a lentiviral library. Transductions were performed at a low multiplicity of infection (MOI ∼0.35), using enough cells to achieve a representation of at least 1,000 transduced cells per sgRNA assuming a 20% - 40% transduction efficiency. The transduction protocol was the same as listed above. Puromycin was added 2 days post-transduction for presence-based selection conditions and 4 days post-transduction for activity-based selection conditions to allow time for editing to occur. Cells were passaged on puromycin for 2-3 passages to ensure complete removal of non-transduced cells. When selection was complete, if drugs were used in the screen, cells were split to any drug arms (each at a representation of at least 1,000 cells per sgRNA) and passaged every 2-4 days for an additional 2 weeks to allow sgRNAs to enrich or deplete; cell counts were taken at each passage to monitor growth. At the conclusion of each screen, cells were pelleted by centrifugation, resuspended in PBS, and frozen promptly for genomic DNA isolation.

### Genomic DNA isolation, PCR, and sequencing

Genomic DNA (gDNA) was isolated using either the KingFisher Flex Purification System with the Mag-Bind Blood & Tissue DNA HDQ Kit (Omega Bio-Tek), or the Macherey Nagel NucleoSpin Blood Maxi (2e7–1e8 cells), Midi (5e6–2e7 cells), or Mini (< 5e6 cells) kits, per the manufacturer’s instructions. The gDNA concentrations were measured by Qubit. For samples where genomic DNA was limited, gDNA was purified prior to PCR using the Zymo OneStep PCR Inhibitor Removal Kit (Zymo), per the manufacturer’s instructions.

For PCR amplification, gDNA was divided into 100 µL reactions such that each well had at most 10 µg of gDNA. Plasmid DNA (pDNA) was also included at a maximum of 100 pg per well. Each well of a 96-well PCR plate contained 1.5 µL of Titanium Taq (Takara), 10 µL of Titanium Taq buffer, 8 µL of dNTPs, 5 µL of DMSO, 0.5 µL of P5 primer at 100 µM stock, 10 µL of P7 primer at 5µM stock, and sterile water added to 100 µL. PCR cycling conditions were as follows: (1) 95°C for 1 min; (2) 94°C for 30 s, (3) 52°C for 30 s, (4) 72°C for 30 s, (5) go to step 1, x28; (6) 72°C for 10 min. PCR products were purified with Agencourt AMPure XP SPRI beads according to manufacturer’s instructions (Beckman Coulter, A63880). Samples were sequenced using Illumina technology (either MiSeq, HiSeq, or NovaSeq) with a 5% spike-in of PhiX.

### Validation experiments

For validation experiments, individual sgRNA vectors were assembled and made into lentivirus as described above. At least 3x10^6^ cells were transduced in duplicate with a virus volume to obtain ∼30% transduction efficiency and were selected with puromycin to remove uninfected cells; puromycin doses and selection timeline were as described above.

For NG-Cas9 base editor validation experiments (related to Figure 1), after puromycin selection was completed, cells were split between no drug and Nutlin conditions and cultured for an additional 14 days. Genomic DNA was isolated using the Kingfisher as described above, and the target sites were amplified using a 2-step PCR. In the first round of PCR, genomic DNA was amplified using custom primers designed to amplify each target site. Each well contained 50 µL of NEBNext High Fidelity 2X PCR Master Mix (New England Biolabs), 0.5 µL of each primer at 100 µM, and 49 µL of gDNA. We used a touchdown PCR with the following cycling conditions: (1) 98°C for 1 min; (2) 98°C for 30 s; (3) 68°C for 30 s (- 1°C per cycle); (4) 72°C for 1 min; (5) Go to step 2, x 15; (6) 72°C for 10 mins. The second round of PCR appended Illumina adapters and well barcodes for sequencing. Each well contained 1.5 µl of Titanium Taq (Takara), 10 µL of Titanium Taq buffer, 8 µL of dNTPs, 5 µL of DMSO, 0.5 µL of P5 primer at 100 µM, 10 µL of P7 primer at 5 µM, 55 µL of water, and 10 µL of PCR product from the first PCR. The following cycling conditions were used: (1) 95°C for 1 min; (2) 94°C for 30 s; (3) 52.5°C for 30 s; (4) 72°C for 30 s; (5) go to step 2, x 15; (6) 72°C for 10 mins. Each well was separately purified with Agencourt AMPure XP SPRI beads according to the manufacturer’s instructions (Beckman Coulter, A63880), using a 1:1 ratio of beads to PCR product. DNA concentration was quantified using a Nanodrop. Sample concentrations were quantified by Qubit. Samples were then sequenced using Illumina MiSeq300 and a 10% PhiX spike-in. FastQ files were then processed with CRISPResso2 to obtain allele-level abundances.

For SpG-Cas9 base editor splice-targeting sgRNA validation experiments, after puromycin selection was completed, MelJuSo cells were passaged with interferon-γ to stimulate *CD274* expression at a concentration of 0.1 µg/mL. Two days later, cells were stained with APC anti-human CD274 antibody (BioLegend 393610). Staining was performed by seeding 150 µL of cells per well in a 96-well U-bottom plate, centrifuging at 1000 x g for 5 min, resuspending in 99 µL flow buffer (PBS +2% FBS, 1% EDTA) and 1 µL antibody. The plate was incubated on ice for 20-30 mins, then cells were washed 3x by centrifugation and resuspension in 200 µL of flow buffer per well. Flow cytometry was performed using a Beckman Coulter CytoFLEX, and the acquired FCS files were analyzed using FlowJo™ Software.

For SpG-Cas9 and SpRY-Cas9 editing efficiency validation experiments, the *CD274* locus was amplified using two custom primers ordered from IDT with sequences as follows: 5’-TTAGAACCACCAAGTCCCAT-3’; 5’-AGAAGACTTTGCCATTGTGT-3’. PCR amplification was performed using the protocol described above, and the resulting amplicons were Sanger sequenced by Azenta. Reads were overlaid and compared with the native CD274 locus using Snapgene.

### Western blot

Cells were prepared for Western blotting by lifting via scraping and lysing with NP-40 lysis buffer (ThermoFisher, FNN002) supplemented with protease inhibitor (Roche, 11836170001). The extracted protein samples were quantified using the Pierce™ BCA Protein Assay kit (ThermoFisher, 23225) to determine loading volume. Samples were loaded at 15-20 µg per well onto a NuPAGE 4-12% Bis-Tris Acrylamide midi gel along with Precision Plus dual color ladder (BioRad, 1610374). The gel was run on an XCell4 Surelock Midi-Cell (ThermoFisher WR0100) with MES buffer (Life Technologies, NP0002) at 120V for 60 mins. The gel was transferred to a membrane using the iBlot2 transfer stack package (ThermoFisher, IB23001). Blocking was done by rocking for 1 h at room temperature in blocking buffer (LI-COR, 927-70050). Primary (Abcam, ab28 (anti-p53) and ab219649 (anti-Vinculin)) and secondary (LI-COR, 925-32210 and 925-68071) antibody solutions were made in blocking buffer supplemented with Tween-20 (VWR, 100216-360). The blot was incubated in the primary antibody solution while rocking at 4°C overnight, followed by 3 x 10 min washes in TBST. Secondary antibody solution was added and the blot was incubated for 1 h at room temperature then washed 3x for 10 min in TBST. The prepared blot was imaged on the LI-COR Odyssey.

### Quantification and statistical analysis

#### Self-editing analysis pipeline

For the SpG-Cas9 NNN-PAM screen, we created a dictionary with all sgRNA sequences as keys and all possible A>G or C>T edits as values. We then took the top 1 million unexpected sequences in terms of total read count frequency and mapped these unexpected sequences to their corresponding original sgRNA sequence had any self-editing occurred. Finally, we added read counts from these unexpected sequences to their mapped sgRNA sequences and proceeded with analysis.

#### Screen analysis

sgRNA sequences were extracted from sequencing reads by running PoolQ with the search prefix “CACCG”, except for the splice-targeting sgRNA screen, which used the prefix “TCCCG”. Reads were counted by alignment to a reference file of all possible sgRNAs present in the library. The read was then assigned to a condition (e.g., a well on the PCR plate) on the basis of the 8 nt index included in the P7 primer. After deconvolution, read counts were log-normalized by the following formula:

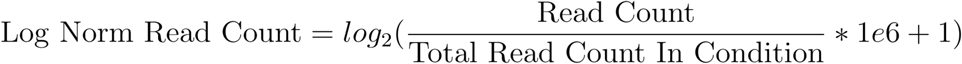

We then calculated the LFCs. All no drug (dropout) conditions were compared to the plasmid DNA (pDNA); drug-treated conditions were compared to the time-matched dropout sample. We assessed the correlation between LFCs of replicates. Prior to further analysis, we filtered out sgRNAs for which the log-normalized reads per million of the pDNA was > 3 standard deviations below the mean. LFCs for the no drug arm were calculated relative to the pDNA. LFCs for the drug arms were calculated relative to the no drug arms. For all pooled screens except the splice-targeting sgRNA screen, z-scores were calculated relative to the intergenic controls. For the splice-targeting sgRNA screen, z-scores were calculated relative to the mean and standard deviation of all replicates.

#### Data visualization

Figures were created with Python and FlowJo™ Software. Schematics were created with BioRender.com.

---

**Supplementary figure 1:**
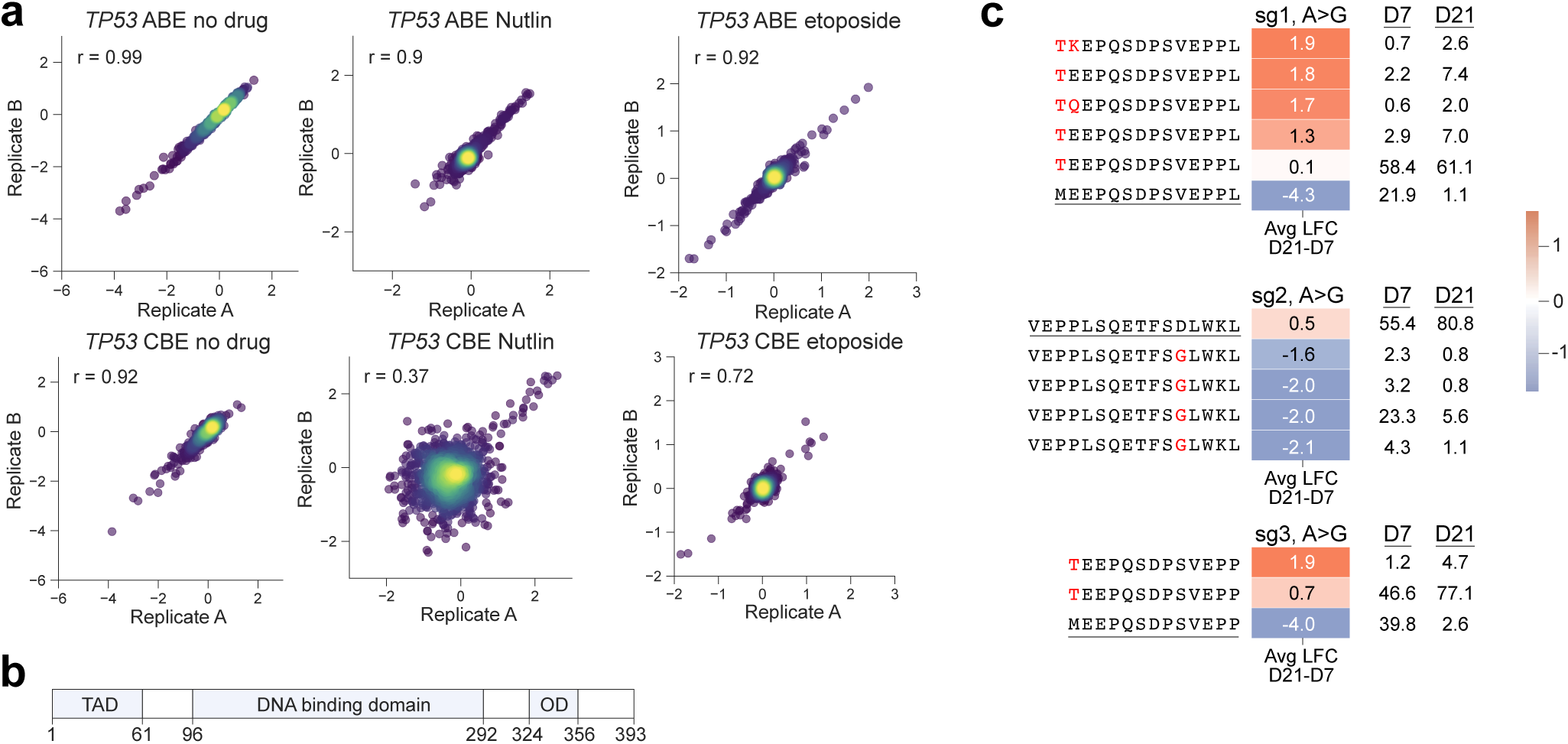
NG-Cas9 base editor screen quality and validation (a) Replicate correlations of ABE (top) and CBE (bottom) arms across all drug conditions. Pearson correlation reported. (b) Annotated TP53 domains. Domain start and end positions are labeled. TAD=transactivation domain, OD=oligomerization domain.(c) TP53 validation target allele abundance of sgRNAs 1, 2, and 3. sgRNAs were transduced into cells expressing NG-Cas9-ABE and treated with Nutlin starting on Day 7. Allelic fractions at each time point were determined by Illumina sequencing. LFC in abundance from Day 7 to Day 21 is shown and color coded. Wildtype sequence is underlined. Mutated amino acids are shown in red.

**Supplementary figure 2:**
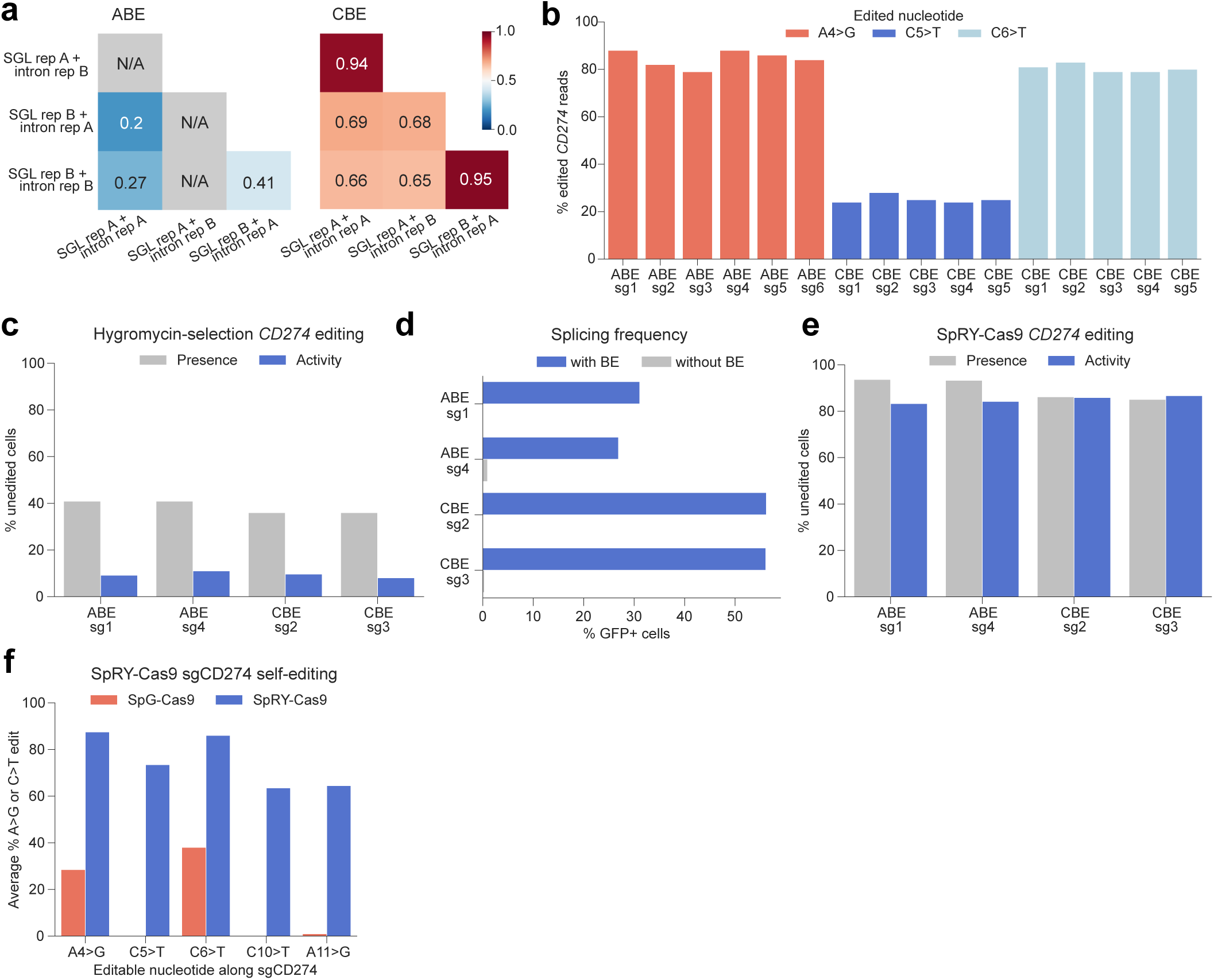
Activity-based selection method development and extension (a) Replicate correlations between splice-targeting guide library (SGL) and intron library replicates for ABEs (left) and CBEs (rightt). Note that the ABE SGL Rep A + Intron Rep B sample was lost to contamination. (b) Percent base editing by nucleotide as determined by Sanger sequencing of targeted CD274 locus. Only within-window edits are shown. (c) Editing efficiency of presence-versus activity-based selection methods with the intron inserted within the hygromycin resistance gene. (d) GFP assay to evaluate possible cryptic splicing by comparing the GFP-intron construct in cells with and without base editors. (e) Editing efficiency of presence- versus activity-based selection methods with puromycin resistance and SpRY-Cas9 base editors. (f) SpRY-Cas9 versus SpG-Cas9 self-editing at the integrated sgRNA locus. Two splice-targeting sgRNAs were used for both ABE and CBE, and the average self-editing is plotted.

**Supplementary figure 3:**
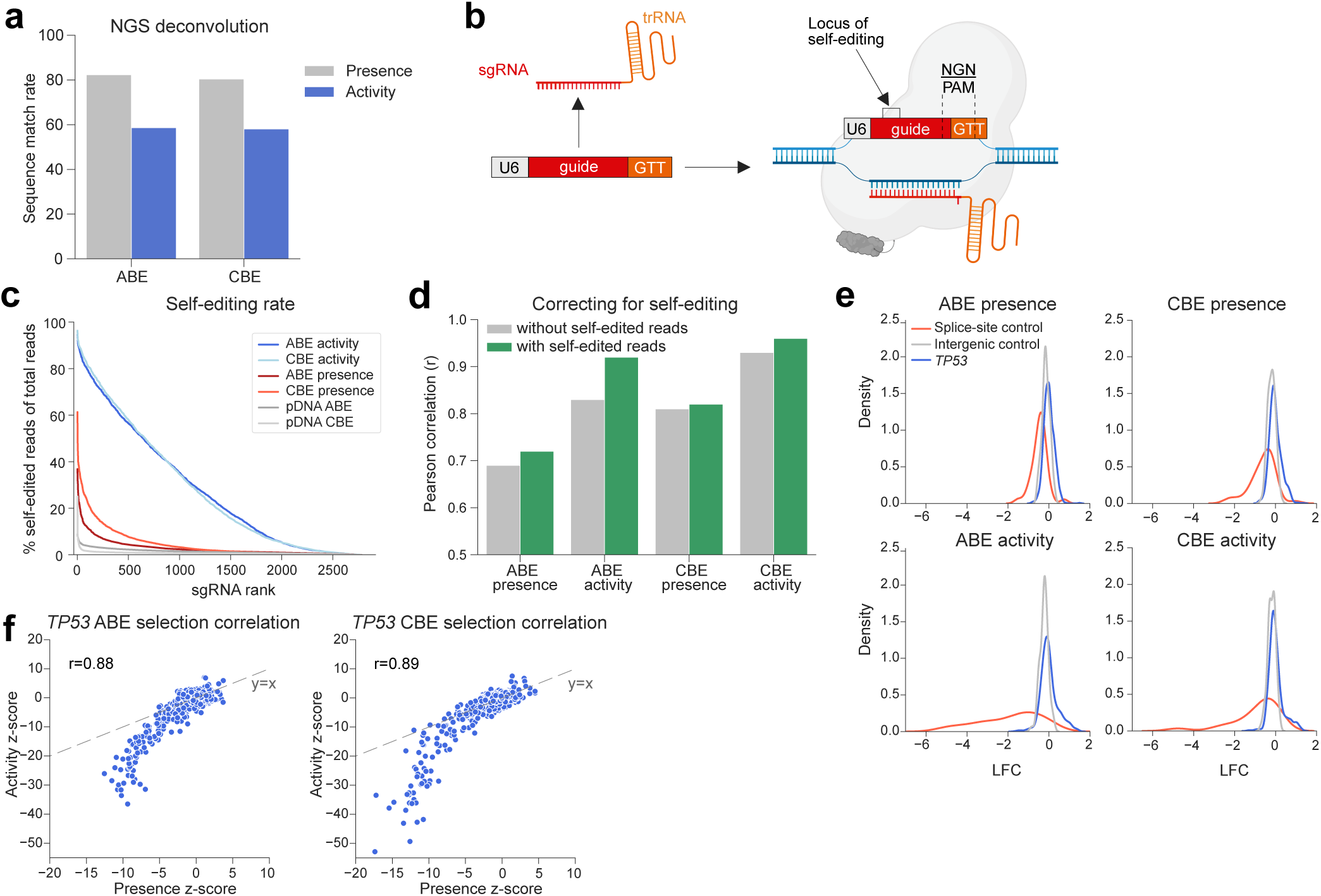
Activity-based selection base editor tiling of TP53 (a) Illumina sequencing read match rate of library sgRNAs after deconvolution with PoolQ. (b) Potential self-editing mechanism. Rather than identifying and editing the endogenous target, the sgRNA may bind to itself by forming a one-nucleotide bulge to recognize the NGN PAM using the first G of the tracer sequence. Alternatively, GTT may function as a PAM sequence. (c) Self-editing rate by selection method and base editor. Shown in gray is baseline pDNA self-editing rate with no editors to account for sequencing error rates. (d) Screen replicate Pearson correlations of the etoposide arm before and after applying self-editing computational correction. (e) sgRNA LFC distribution graphs for ABE (left) and CBE (right) with presence-based selection (top) and activity-based selection (bottom) conditions, comparing the no drug final time point to pDNA sequencing. Splice-site control sgRNAs target pan-lethal genes. (f) sgRNA distribution between selection conditions for ABE (left) and CBE (right) arms. Pearson correlations are displayed. Dashed line indicates y = x.

